# A Novel Mouse Model of Parkinson’s Disease for Investigating Progressive Pathology and Neuroprotection

**DOI:** 10.1101/2025.02.13.638053

**Authors:** Santhosh Kumar Subramanya, Shreshth Shekhar, Devu B Kumar, Sathyapriya Senthil, Dharmaraja Allimuthu, Kelvin C Luk, Poonam Thakur

## Abstract

Developing animal models that successfully recapitulate the features of progressive Parkinson’s disease (PD) is crucial for understanding disease progression mechanisms and creating effective therapeutic interventions. In this study, we created a mouse model of PD by overexpressing α-synuclein through a combined injection of AAV6-α-synuclein and preformed fibrils (PFFs) into the medial and lateral substantia nigra (SN). We also demonstrated that chronic administration of the c-Abl inhibitor PD180970 provides neuroprotection in this model.

Mice injected with the AAV6-α-synuclein and PFF combination showed a progressive loss of dopaminergic (DA) neurons in the SN and their projections in the striatum over 24 weeks. This neuronal loss coincided with a time-dependent accumulation of phosphorylated α-synuclein (p-syn) in the SN. The p-syn aggregates spread to synaptically connected DARPP-32-positive neurons in the striatum and further extended to the cortex. We also observed a contralateral spread of p-syn aggregates. Additionally, α-synuclein overexpression led to a significant increase in activated microglia and astrocytes at all timepoints, with the strongest activation occurring early and gradually diminishing over time.

Daily administration of PD180970 significantly reduced the loss of DA neurons caused by α-synuclein injection and decreased the accumulation of p-syn in the SN. PD180970 treatment also reduced the neuroinflammation significantly.

Overall, the combined injection of AAV6-α-synuclein and preformed fibrils into the mouse brain establishes a robust PD model, enabling detailed mechanistic studies of the disease. We further demonstrate the model’s utility for chronic neuroprotection studies using the potential drug PD180970, highlighting its broad applicability.

**Significance Statement:** This study establishes a robust mouse model of Parkinson’s disease (PD) by combining AAV6-mediated α-synuclein overexpression and preformed fibrils (PFFs) to replicate key features of PD, such as progressive dopaminergic neuron loss, phosphorylated α-synuclein accumulation, and neuroinflammation. The model captures the spread of pathological aggregates to synaptically connected brain regions, closely mimicking the human disease. By testing the c-Abl inhibitor PD180970, we demonstrate its neuroprotective effects, including reduced neuronal loss, decreased α-synuclein accumulation, and neuroinflammation highlighting its therapeutic potential. This model offers a valuable platform for investigating PD mechanisms and evaluating novel interventions, bridging the gap between preclinical and clinical applications.

## Introduction

Parkinson’s disease (PD) is an age-related neurodegenerative disorder marked by the loss of dopaminergic (DA) neurons in the substantia nigra (SN), leading to motor deficits (1). This neuronal loss is driven by multiple pathological processes such as abnormal protein aggregation, neuroinflammation, impaired protein quality control, and organelle dyshomeostasis, among others (2). Despite advances in understanding the mechanisms underlying PD, there is no cure for this disease, and current therapies are primarily focused on managing symptoms.

Animal models are crucial for advancing our understanding of PD pathogenesis and preclinical evaluation of potential therapeutics. Traditional toxin-based models such as MPTP and 6-OHDA (3–9) induce acute DA neuron loss and do not replicate the slow, multifaceted neurodegeneration characteristic of PD (10, 11). Transgenic models such as SNCA mutants and LRRK2 mutants produce mild phenotypes that typically appear only in aged mice (12–18). Although these models work well for studying prodromal PD, their prolonged timelines and high costs often make them impractical for neuroprotection studies. Chronic PD models induced by intranigral or intrastriatal injection of adeno-associated virus (AAV)-α-synuclein present a balance between these two as they cause progressive neurodegeneration within shorter timelines compared to transgenic models (19–21). However, these models also come with limitations. High viral titers are often required to achieve sufficient α-synuclein overexpression for neurodegeneration, but such titers can trigger non-specific cellular toxicity unrelated to α-synuclein expression (22, 23). Further, the resulting α-synuclein levels often exceed the physiological range (23). Conversely, low titers may not induce sufficient expression to drive the neurodegenerative process. The discovery of fibrillar α-synuclein in PD brains has spurred the development of models using pre-formed fibrils (PFF) generated from recombinant α-synuclein protein (24–26). These models demonstrate neuronal loss and pathological features over a slow timescale of 9-12 months (25–27). Combining PFFs with AAV-α-synuclein allows for the utilization of lower viral titers, minimizing generic cell toxicity (28, 29). This approach leverages the templated seeding principle, where PFFs act as aggregation nuclei for AAV-driven α-synuclein overexpression. Since AAVs cause the sustained expression of α-synuclein, the model is also progressive in nature (28). This combinatorial model has been termed the SynFib model (29).

The SynFib model has demonstrated efficacy in rats, reproducing disease features such as motor impairments, neuroinflammation, and α-synuclein aggregation (22, 28–31). However, its applicability in mice remains poorly understood. Given the widespread use of mice in research, adapting this model for mice could offer a valuable tool for the broader scientific community. In this study, we developed and characterized a SynFib model in mice using an AAV-expressing human wild-type α-synuclein combined with PFFs generated from full-length human α-synuclein, creating a humanized model of PD. While some rat models require sequential injections of AAV-α-synuclein and PFF, necessitating two surgeries, we used a single surgery for simultaneous injection of AAV-α-synuclein and PFF in mice SN.

This approach removes the need for a second surgery, thereby reducing animal distress and mortality risk. To ensure coverage of the entire nigra, we performed the injection in both the medial and lateral SN regions. We assessed mice using a battery of motor tests and immunohistochemical (IHC) analyses to evaluate motor deficits, neurodegeneration, protein aggregation, the spread of aggregates in various brain regions, and neuroinflammation.

To validate the utility of this model, we evaluated the neuroprotective effects of PD180970, an orthosteric inhibitor of c-Abl (32). c-Abl upregulation has been observed in the postmortem striatal samples of PD patients (21, 33–35). Moreover, c-Abl depletion or knockout has conferred resistance to PD pathophysiology in various models, including MPTP, α-synuclein PFF, and A53T transgenic mice (34, 36, 37). Due to the pleiotropic effects of c-Abl in cellular physiology, targeting it for neuroprotection offers an ideal opportunity to simultaneously target multiple pathophysiological processes in PD. PD180970 has shown neuroprotective effects in an acute MPTP mouse model by promoting autophagic clearance of α-synuclein aggregates and reducing neuroinflammation while also improving motor deficits

(38). However, to the best of our knowledge, no studies have assessed the safety and efficacy of PD180970 in a chronic model of PD that faithfully recapitulates disease progression. Further, previous studies targeting c-Abl in PD used only male animals (39, 40) while we examined the effects of PD180970 in both sexes for the first time.

## Results

### SynFib group displayed progressive loss of DA neurons and motor deficits

Behavioral and histological analysis were conducted to evaluate the effects of GFP, Fib, Syn, and SynFib exposure on motor activity and neurodegeneration (Fig 1A). At 4W, mice in all groups showed comparable locomotor performance in the wire hang test (Fig S1A), gait analysis (Fig S1B), cylinder test (Fig S1C) and open field test (Fig S1D). Although Syn mice exhibited a shorter latency to fall in comparison to the other groups in the wire hang test the difference was not statistically significant (Fig S1A).

**Fig 1:**
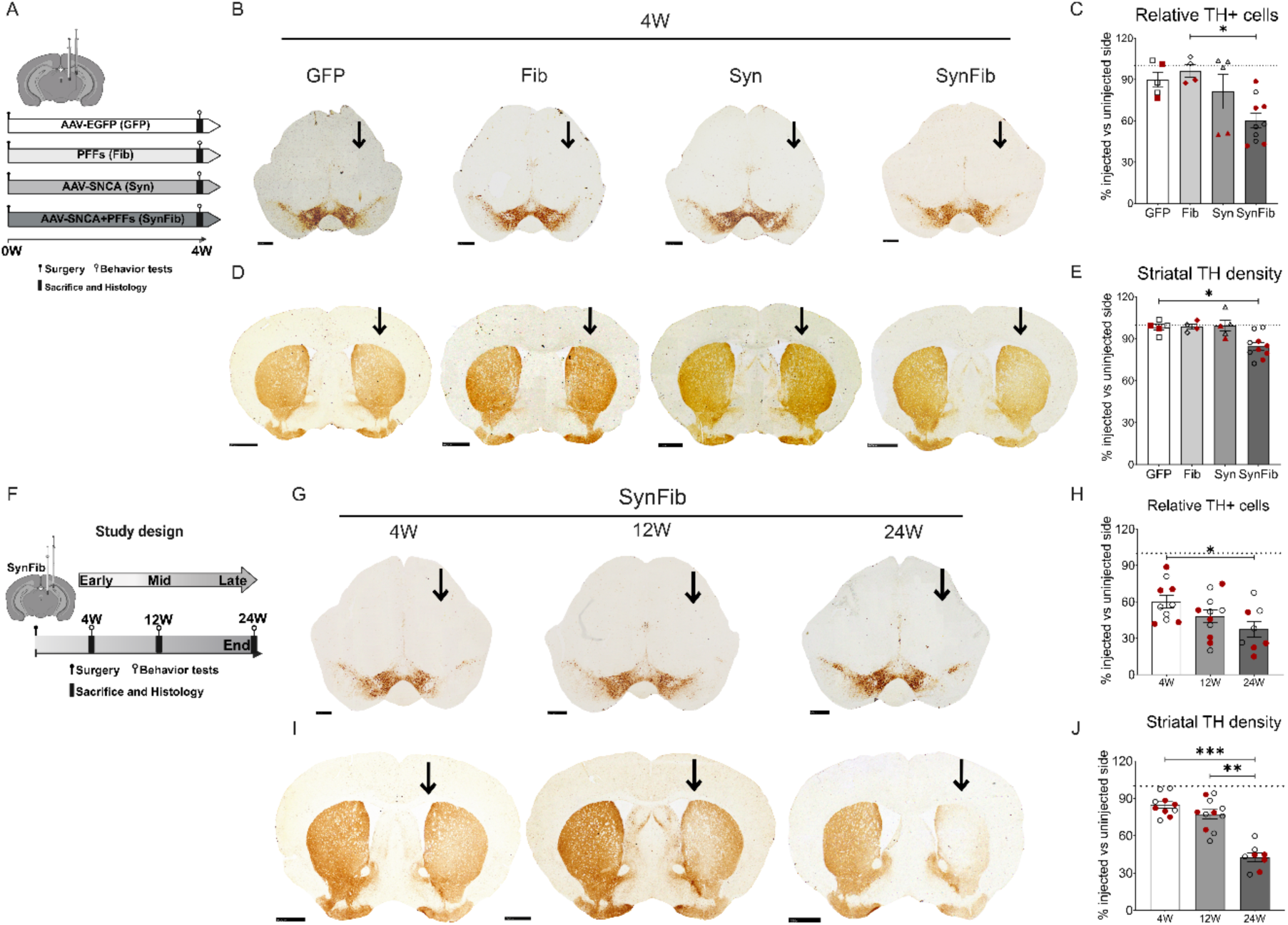
The SynFib model displays progressive neurodegeneration and gait deficits with time. **A.** Experimental design: Mice were stereotaxically injected with AAV6-EGFP (GFP), PFFs (Fib), AAV6-SNCA (Syn), or a combination of AAV6-SNCA and PFF (SynFib) and sacrificed after 4W. **B.** Representative TH-stained sections of the SN in GFP, Fib, Syn, and SynFib groups. Vertical arrow denotes the injected side. Scale bar = 500 µm. **C.** Quantification of TH-positive cells in the injected side relative to the uninjected side. The SynFib group showed significant TH cell loss compared to the Fib group. No of mice: GFP (n = 5), Fib (n = 4), Syn (n = 5), SynFib (n = 10). **D.** Representative TH-stained sections of the STR in GFP, Fib, Syn, and SynFib groups;Scale bar = 1000 µm**. E.** Relative TH fiber density in the STR revealed significant loss in the SynFib group compared to the GFP group. No of mice: GFP (n = 5), Fib (n = 4), Syn (n = 5), SynFib (n = 10). **F.** Experimental timeline for disease progression in the SynFib model at 4W (early), 12W (mid), and 24W (late) stages. **G.** Representative TH-stained sections of the SN in the SynFib group at 4W, 12W, and 24W. Vertical arrow denotes the injected side. Scale bar = 500 µm. **H.** Quantification of TH-positive cells in the injected side at 4W, 12W, and 24W, showing a significant decrease over time. No of mice: 4W (n = 10), 12W (n = 11), 24W (n = 8). **I.** Representative TH-stained sections of the STR in the SynFib group at 4W, 12W, and 24W. Vertical arrow denotes the injected side. Scale bar = 1000 µm. **J.** Relative TH fiber density in the STR at 4W, 12W, and 24W, revealing a significant decrease over time. No of mice: 4W (n = 10), 12W (n = 11), 24W (n = 8). All data are represented as mean ± SEM. Statistical analysis was performed using one-way ANOVA followed by Kruskal-Wallis multiple comparison post hoc test (*P < 0.05, **P < 0.01, ***P < 0.001). Data points in red represent female mice; black represents male mice.

AAV-GFP-injected mice showed only a marginal decline (10% w.r.t. to uninjected side) in TH-positive DA neurons in the SN (Fig 1B and C), suggesting no significant general toxicity from the AAV injection or the titer used in this study. Similarly, mice injected with AAV-synuclein exhibited minimal cell loss (19%), confirming that the α-synuclein expression achieved with low AAV titers is insufficient to cause substantial neurodegeneration at early timepoints. Injection of PFFs alone also resulted in negligible TH neurodegeneration (4%) (Fig 1C). In contrast, SynFib group mice displayed approximately 40% loss of TH-positive neurons at 4 weeks (Fig 1C), along with a 15% reduction in TH density in the striatum (Fig 1D and E). These results demonstrate that the combination of PFFs and AAV-α-synuclein induces significant cell loss at 4W, which is not achieved by either of them individually.

To determine whether the motor deficits and neurodegeneration observed in the SynFib group is progressive, we extended our study to 12W and 24W post-surgery (Fig 1F). We observed no significant decline in the wire hang test (Fig S1E). However, mice exhibited a significant decrease in average stride length at 24W compared to 4W, indicating gait abnormalities (Fig S1F). There was no significant change in the cylinder test (Fig S1G). Mice also showed a significant reduction in time spent in the inner zone of the open field at 12W in comparison to 4W, suggesting anxious behavior (Fig S1H).

Further, SynFib mice exhibited additional SN DA neuron loss (52%) at 12W (Fig 1G and H). This gradual decline continued, reaching 62% at 24W (significant compared to the 4W timepoint) (Fig 1G and H). A similar time-dependent reduction in striatal TH density in the injected side in comparison to the uninjected side was observed, with losses of 15% at 4W, 23% at 12W, and 58% at 24W (Fig 1I and J). These findings demonstrate that the SynFib model exhibits progressive, albeit non-linear, DA neuron degeneration.

To determine whether the observed TH cell loss reflects neuronal death or merely TH downregulation, we performed co-immunostaining of TH with HuD+HuC, a general neuronal marker, using single slices from each animal at all three timepoints (Fig S2A-C). Quantification revealed approximately 17% loss of HuD+HuC-positive cells at 4W, 41% at 12W, and 65% at 24W in the SN (Figure S2D). These results suggest that the observed DA cell loss can be partially attributed to TH downregulation at all timepoints.

### Increased accumulation of α-synuclein aggregates in the nigrostriatal system of the SynFib group

Phosphorylation of α-synuclein at Ser129 is a key post-translational modification linked to its aggregation. To assess α-synuclein aggregation, p-syn immunostaining was performed across all groups. The SynFib (∼2.7-fold higher than GFP) and Syn (∼2.2-fold higher than GFP) groups exhibited elevated p-syn levels on the injected side (Fig 2A and B). In contrast, no aggregates were observed in the GFP group (Fig 2A and B) and the Fib group displayed only a modest 1.2-fold increase in p-syn levels relative to the GFP group. These observations suggest that overexpression of α-synuclein drives the appearance of p-syn aggregates. This observation is consistent with previous studies indicating that PFFs require extended periods for aggregate formation and spread (16). Notably, only the SynFib group displayed striatal p-syn accumulation at 4W (Fig 2C), indicating that aggregates had travelled along the nigrostriatal pathway, successfully replicating pathological spread.

**Fig 2:**
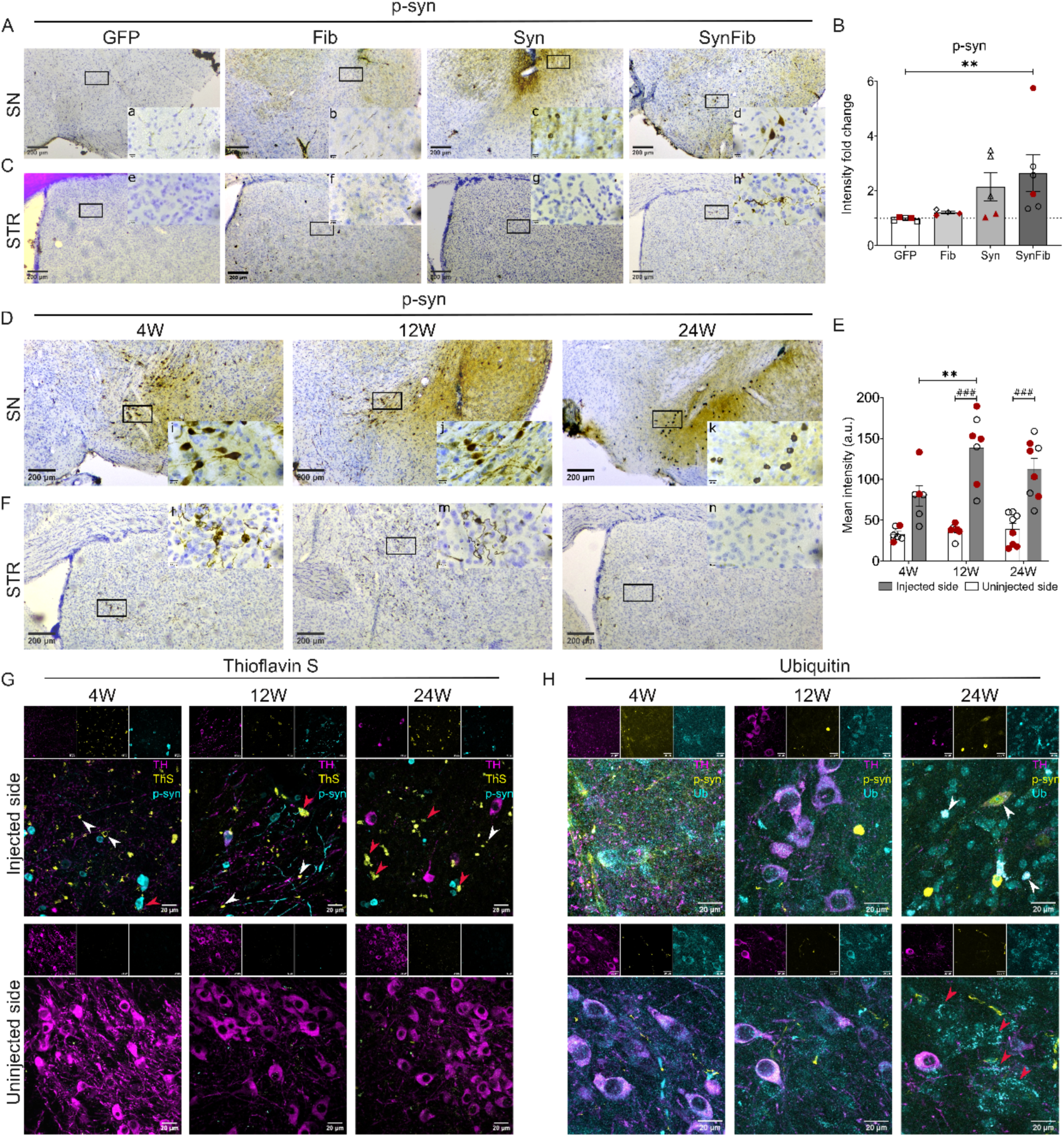
SynFib model displays a high accumulation of p-syn in nigro-striatal pathway. **A.** Representative SN sections stained for p-syn in GFP, Fib, Syn, and SynFib groups at 4W. p-syn-positive aggregates were observed in the Syn and SynFib groups. Scale bar = 200 µm (low magnification) and 20 µm (high magnification). **B.** Relative p-syn levels in SN of GFP, Fib, Syn and SynFib groups. A significantly increased p-syn signal in the SynFib group relative to the GFP group was observed. A non-significant increase in the Syn group was also observed. No of mice: GFP (n = 5), Fib (n = 4), Syn (n = 5), SynFib (n = 6). All data are represented as mean ± SEM. Non-parametric one-way ANOVA followed by Kruskal-Wallis multiple comparison post hoc test (** P < 0.01). Data points in red represent female mice; black represents male mice. **C.** Representative STR sections stained for p-syn in GFP, Fib, Syn and SynFib groups. p-syn aggregates were observed only in the SynFib group. Scale bar = 200 µm (low magnification) and 20 µm (high magnification). **D.** Representative SN sections stained for p-syn in the SynFib group at 4, 12 and 24W. High-magnification images i-k illustrate morphological changes: i-diffuse p-syn in cell body and thin axons, j-axonal swelling filled with p-syn aggregates giving a beaded appearance and k-dystrophic cells with no processes. Scale bar = 200 µm (low magnification) and 20 µm (high magnification). **E.** Mean intensity of p-syn staining in uninjected and injected SN at 4, 12 and 24W. A significant increase in the injected side relative to uninjected is observed at 12 and 24W. A significant increase in abundance of p-syn levels with time (12W relative to 4W) was also observed. No of mice: 4W (n = 6), 12W (n = 7), 24W (n = 8). All data are represented as mean ± SEM. Two-way ANOVA with Tukey’s multiple comparison as post hoc. (### P < 0.001, ** P < 0.01). * indicates comparison between timepoints; # indicates comparison between injected and uninjected side. Data points in red represent female mice; black represents male mice. **F.** Representative STR sections stained for p-syn in the SynFib group at 4, 12 and 24W. High magnification insets l-n reveal axonal thickening and neurites at 4 and 12W but their lack at 24W. Scale bar = 200 µm (low magnification) and 20 µm (high magnification). **G.** Representative SN sections co-stained for p-syn, ThS and TH at 4, 12 and 24W. Top panel - p-syn-positive neurons in the SN co-stained with ThS (red arrowheads) in the SynFib group, confirming the amyloid nature of aggregates. Some ThS positive deposits were p-syn negative (white arrowheads) at 4-24W depicting non-phosphorylated aggregates. Bottom panel-No ThS positive deposits were observed in the uninjected side. Scale bar = 20µm. **H.** Representative SN sections from SynFib mice co-stained for p-syn, ubiquitin and TH at 4, 12 and 24W. Top panel-Neurons in the injected side show abundant ubiquitin puncta at all timepoints. At 24W, larger, clustered ubiquitin deposits were observed on the injected side (white arrowheads). Ubiquitin puncta were absent on the uninjected side at 4W and 12W but prominent at 24W (red arrowheads). Scale bar = 20µm.

We also investigated the p-syn levels in the SynFib group across 4-24W. Data showed a significant increase in the p-syn positive aggregates in the SN of the injected side compared to the uninjected side at all time points (except 4W, which was non-significant) (Fig 2D and E). P-syn accumulation increased from 4W to 12W, suggesting progressive aggregate buildup. However, a mild decrease in p-syn levels at 24W compared to 12W indicates the loss of cells containing these aggregates (Fig 2E). At 4W and 12W, p-syn accumulation was observed in the cytoplasm, nucleus, and dendrites of the neurons (Fig 2Di,j). Swelling of neuronal processes with p-syn aggregates was particularly prominent at 12W (Fig 2Dj). By 24W, neurons with p-syn aggregates lacked projections and typical neuronal morphology, suggesting their atrophy (Fig 2Dk). In the STR, p-syn-positive neurons and processes were prominent at 4W and 12W but relatively sparse at 24W (Fig 2F). Furthermore, the colocalization of TH with p-syn signal was higher at 4W on the injected side compared to the uninjected side but colocalization was decreased over time (Fig. S3A). This decline may be attributed to the degeneration of p-syn-positive neurons and, in part, to TH downregulation, suggesting the progressive loss of DA neurons bearing p-syn aggregates.

### SynFib group displayed lewy-like pathological features

To further characterize the aggregates, we stained SN sections with Thioflavin-S (ThS) dye that specifically binds amyloid protein structures. GFP, Fib, and Syn groups showed no ThS-positive aggregates (Fig S3B). In contrast, ThS staining was observed in the SynFib group at all the timepoints (Fig 2G). While some ThS-positive aggregates co-localized with the p-syn signal (Fig 2G, upper panel, red arrowheads), several did not (white arrowheads), suggesting that p-syn staining captures only a subset of pathological aggregates. Additionally, SN sections from 4W, 12W, and 24W SynFib mice were treated with proteinase-K and stained for p-syn. All timepoints displayed p-syn-positive structures resistant to proteinase-K digestion, indicating the presence of mature, insoluble aggregates (Fig S3C).

We also observed changes in ubiquitination patterns with disease progression. At 4W, ubiquitin deposits colocalizing with p-syn appeared as punctate structures throughout the cytoplasm (Fig 2H, top panel). By 24W, ubiquitin was predominantly localized in the nucleus (Fig 2H). Interestingly, changes were also noted on the uninjected side of the SN (Fig 2H, bottom panel). At 4W and 12W, a diffuse ubiquitin signal was observed in the cytoplasm, but by 24W, as p-syn pathology spread contralaterally, punctate ubiquitin structures emerged, suggesting neurons may be attempting to clear aggregates via the ubiquitin-proteasome system (UPS) (Fig 2H).

### Transmission of α-synuclein aggregates beyond DA neurons

Next, we explored the trans-synaptic spread of aggregates. The DA neurons from the SN form synapses with GABAergic medium spiny neurons (MSNs) in the STR (41). Thus, we co-immunostained p-syn with TH and DARPP-32, a marker of MSNs. We observed p-syn aggregates within MSNs at 4W and 12W, suggesting trans-synaptic spread (Fig 3A). However, no co-localization was observed at 24W (Fig 3A) potentially due to the death of aggregate-bearing cells or their clearance by cellular machinery. To verify these possibilities, we quantified the relative fluorescence intensity of DARPP-32 in the STR across 4-24W. Despite the presence of p-syn, DARPP-32 levels on the injected side remained consistent over time (Fig 3B and C), indicating that MSNs can clear aggregates over time, unlike SN DA neurons, which degenerate. Interestingly, p-syn positive aggregates were also found in cortical regions at 4W and 12W but were absent at 24W (Fig 3D), suggesting cortical neurons can effectively handle aggregates.

**Fig 3:**
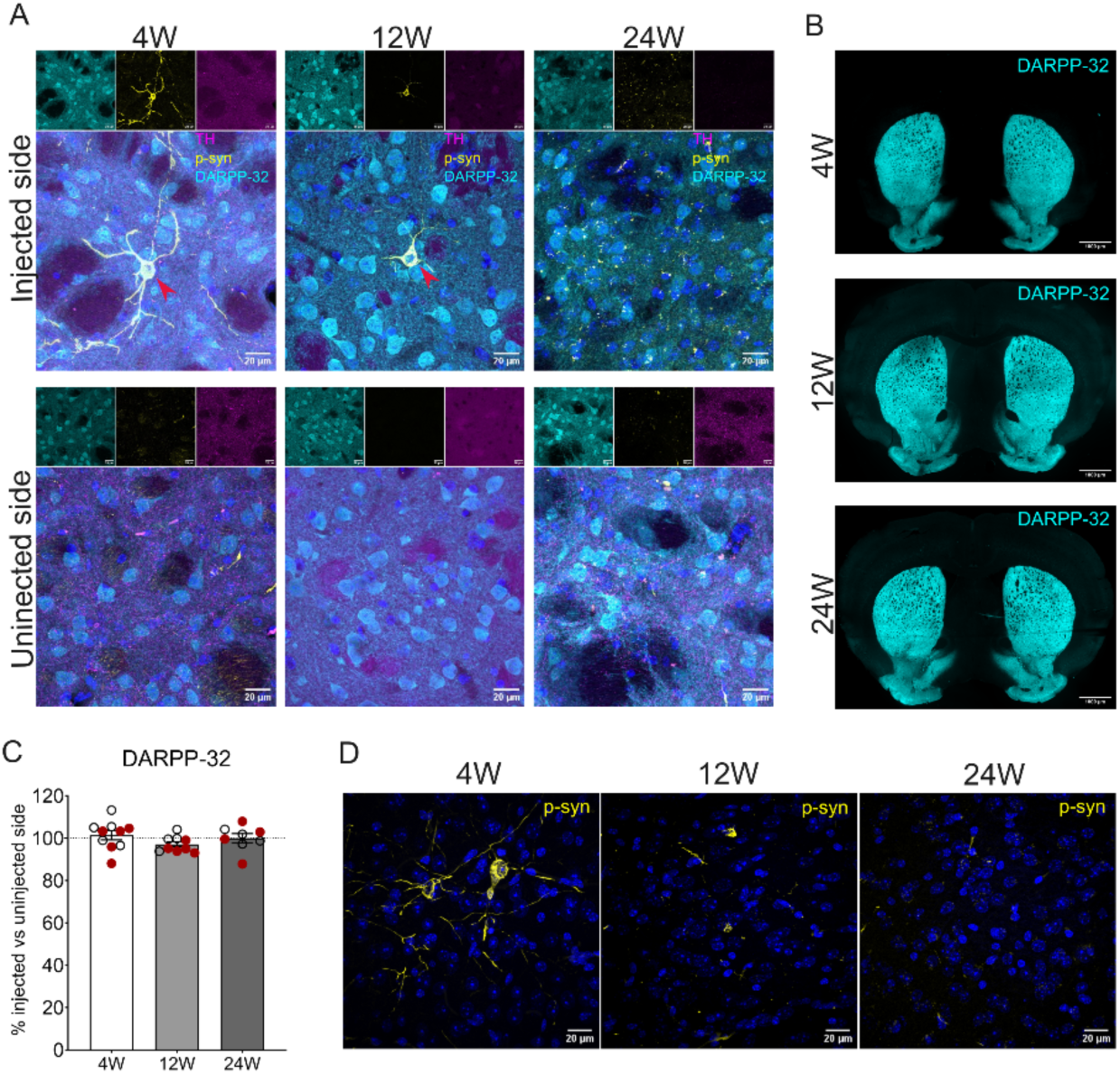
α-Synuclein aggregates spread to MSNs in the striatum and further to the cortex. **A.** Representative high-magnification images of the STR in the SynFib group co-stained for DARPP-32, TH and p-syn at 4, 12 and 24W. Co-localization of p-syn with MSNs in STR demonstrates trans-synaptic spread (Red arrow heads). Scale bar = 20µm. **B.** Representative STR sections from SynFib group stained for DARPP-32 at 4, 12 and 24W. Scale bar = 1000µm **C.** Quantification of relative DARPP-32 intensity at 4, 12 and 24W indicates no significant loss of staining intensity over time suggesting resistance to p-syn pathology. No of mice: 4W (n = 10), 12W (n = 9), 24W (n = 8). All data are represented as mean ± SEM. Statistical analysis was performed using one-way ANOVA followed by Kruskal-Wallis multiple comparison post hoc test. Data points in red represent female mice; black represents male mice. **D.** Representative high-magnification images of p-syn positive structures in the cortex at 4, 12 and 24W in the SynFib group showing their spread beyond the dopaminergic system. Scale bar = 20µm.

### The SynFib model displayed elevation of neuro-inflammation at the early timepoint

PD is associated with neuroinflammation, marked by the activation of microglia and astrocytes. We observed needle tracks with local accumulation of microglia (IBA1) and astrocytes (GFAP) in SN sections from all the groups (Fig 4A and B) at 4W. However, only SynFib group had a significant increase in the size and thickness of the glial processes indicated by upregulation of the area occupied by IBA1 and GFAP in the injected SN in comparison to the uninjected side (Fig 4C and D). We further examined neuroinflammation changes in the SynFib group over time (Fig 4 E and F), observing a time-dependent decline in the area occupied by microglia and astrocytes in the SN (Fig 4G and H).

**Fig 4:**
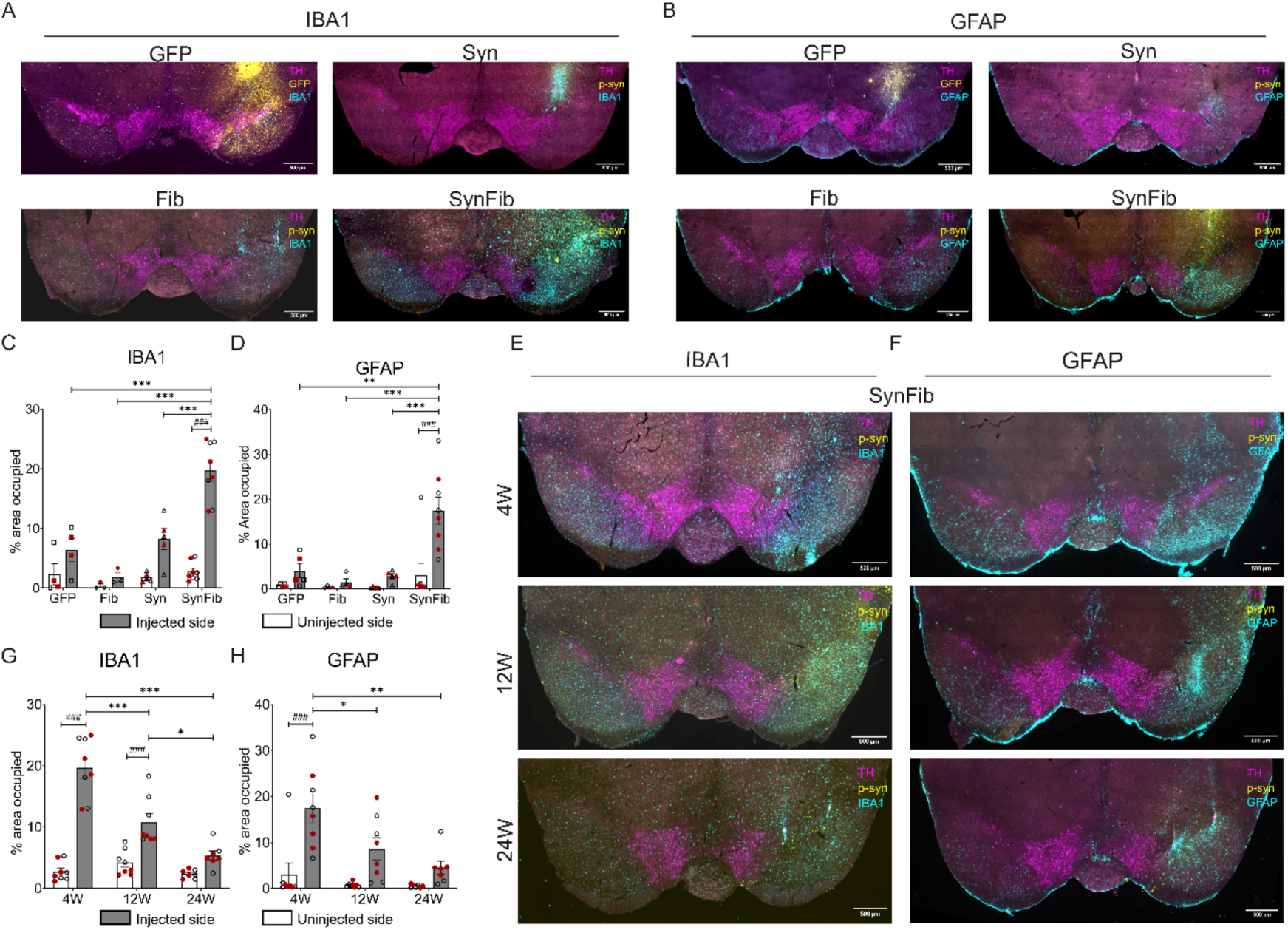
Enhanced neuroinflammation is observed in the SynFib group. **A.** Representative SN sections co-stained for IBA1, p-syn and TH in GFP, Fib, Syn, and SynFib groups at 4W. GFP group shows expression of GFP (yellow). Scale bars = 500µm**. B.** Representative SN sections co-stained for GFAP, p-syn and TH in GFP, Fib, Syn, and SynFib groups at 4W. GFP group shows expression of GFP (yellow). Scale bars = 500µm. **C.** Quantification of the percentage area occupied by IBA1 and **D.** GFAP in GFP, Fib, Syn, and SynFib groups revealed their activation. No of mice for IBA1: GFP (n = 4), Fib (n = 3), Syn (n = 5), SynFib (n = 8) and No. of mice for GFAP: GFP (n = 5), Fib (n = 4), Syn (n = 5), SynFib (n = 8). **E.** Representative SN sections of IBA1 and **F.** GFAP co-stained p-syn and TH in the SynFib group at 4W, 12W, and 24W. Scale bars = 500µm. **G.** Quantification of the percentage of the area occupied by IBA1 and **H.** GFAP in SynFib group across 4W, 12W, and 24W, revealing increased neuroinflammation on the injected side compared to the uninjected side and a time-dependent reduction in area occupied by IBA1 and GFAP. No of mice for IBA1: 4W (n = 8), 12W (n = 8), 24W (n = 7); No of mice for GFAP: 4W (n = 8), 12W (n = 8), 24W (n = 7). Data are represented as mean ±SEM. (* comparison between injected sides across groups; * P<0.05, ** P <0.01, *** P <0.001) (# comparison between uninjected and injected sides;;### P <0.001). Two-way ANOVA with Tukey’s multiple comparison as post hoc. Data points in red represent female mice; black represents male mice.

A closer look at the microglial and astrocytic morphology, at higher magnification revealed that p-syn signal often colocalized with IBA1-positive cells with a (Fig S4A, upper panel). We observed a non-significant increase in colocalization of p-syn with IBA1 in injected side compared to the uninjected side at 4W. This trend reduces with time (Fig S4C, upper panel) potentially due to loss of p-syn cells. Some of the p-syn positive aggregates were also observed inside the microglia on the uninjected side (Fig S4A, lower panel, red arrowheads). Further, we quantified colocalization of p-syn with GFAP positive signals. Although, a non-significant increase was observed in colocalization of p-syn with GFAP at 4W, there were no obvious changes at 12 and 24W between injected and uninjected sides (Fig S4C, lower panel).

### Administration of PD180970 to SynFib model confers neuroprotection

Finally, we explored the utility of the SynFib model for in-vivo evaluation of therapeutic molecules. Studies have shown activation of c-Abl, a non-receptor tyrosine kinase, in both PD brains and animal models. We selected PD180970, a known c-Abl inhibitor, for our experiments. To assess its specificity, we conducted an off-target screen using the Swiss Target Prediction web tool. Consistent with previous reports, PD180970 showed higher selectivity for the non-receptor tyrosine kinase c-Abl1, ranking among the top three predicted targets, over receptor tyrosine kinases such as c-Kit, PDGFRα, and PDGFRβ (Fig S5A). To further investigate PD180970’s binding to c-Abl, we performed molecular docking using Autodock software for both active and inactive c-Abl conformations. PD180970 bound to both conformations, with stronger binding energy for the active form (−14.40 kcal/mol) compared to the inactive form (−9.77 kcal/mol) (Figure S5B and C).

SynFib mice were administered PD180970 for 16W (Fig 5A). Behavioral evaluations revealed no significant differences in motor performance across treatment groups (Fig 5B). Histological analysis of TH-stained sections showed a significant reduction in TH-positive cells in the SN of the SynFib group compared to controls (Fig 5C and D). However, PD180970 administration provided significant neuroprotection to the SynFib mice (Fig 5C and D). This effect was also reflected in striatal TH density, where drug treatment mitigated TH loss (Fig 5E and F). Notably, while both male and female mice exhibited similar neuronal loss, the neuroprotective effects of the drug were more pronounced in females. Additionally, PD180970 significantly reduced p-syn levels in the SynFib group (Fig 5G and H).

**Fig 5:**
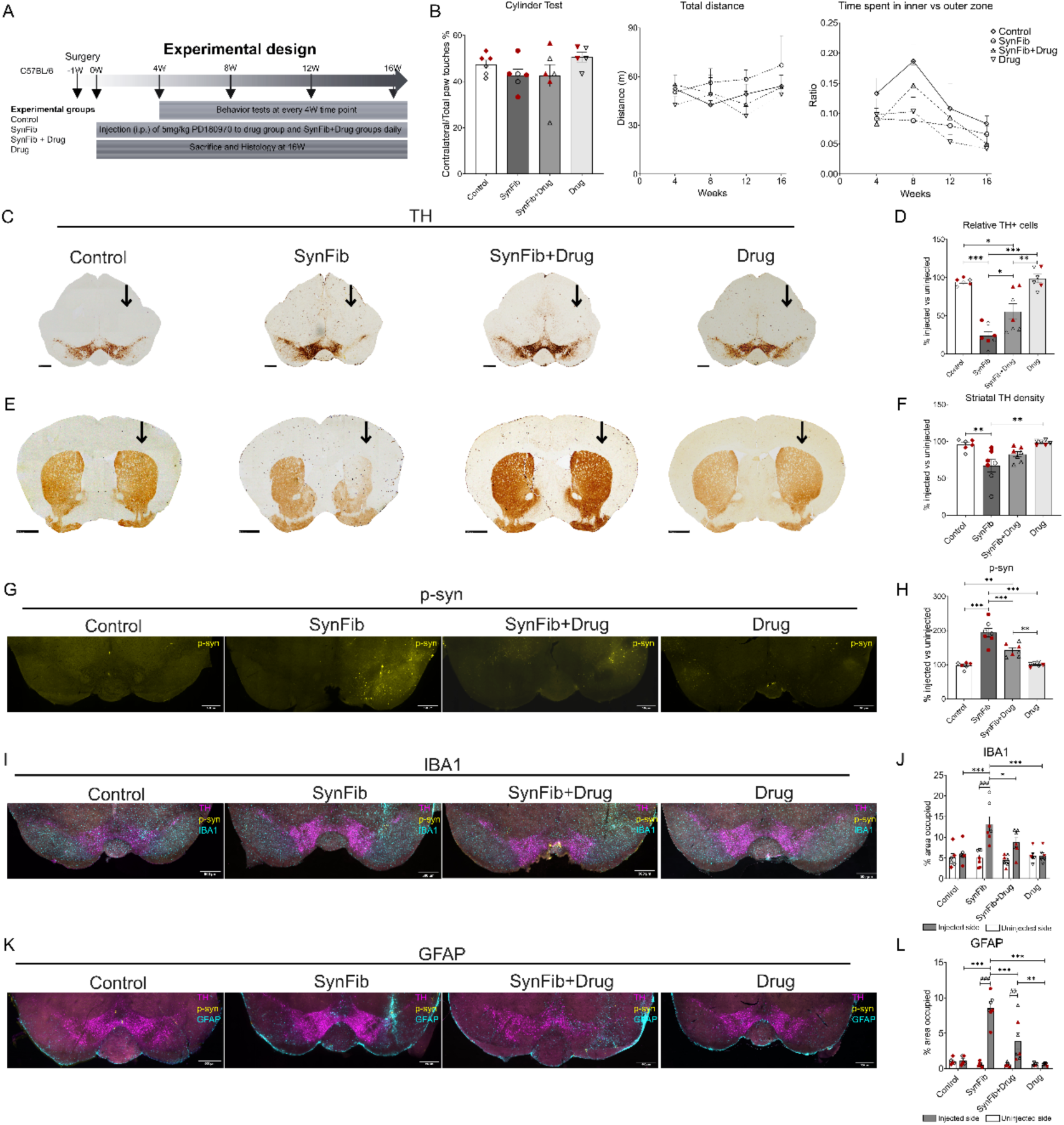
PD180970 administration provides neuroprotection in the SynFib model. **A.** Experimental plan depicting timelines of Control, SynFib, SynFib+Drug and Drug treatment groups. **B.** Behavioral tests: Cylinder test (left panel)-A modest (non-significant) reduction in contralateral paw usage was observed in the SynFib group, which remained unchanged after drug treatment. Open field test (middle and right panel)—The SynFib group exhibited increased total distance traveled (non-significant relative to control), which was attenuated upon drug administration (non-significant relative to SynFib). No significant differences in the time spent in the inner vs. outer zone of the open field were observed between groups. Data are represented as mean + SEM. Control (n=6), SynFib (n=6), SynFib+Drug (n=6), Drug (n=5) for cylinder and Control (n=6), SynFib (n=7), SynFib+Drug (n=7), Drug (n=6) for open field test). Ordinary one-way ANOVA for cylinder test, and two-way ANOVA with Tukey’s multiple comparison test as the post hoc analysis for open field tests. **C** Representative images of SN stained for TH illustrate neuroprotection in the SynFib+Drug group (vertical arrow depicts injected side), Scale bar = 500µm. **D.** Quantitative analysis of relative cell counts in the SN depicting neuroprotection by drug treatment. **E.** Representative images of STR stained for TH, Scale bar = 1000µm **F.** Quantitative analysis of relative TH fiber density in the STR depicting neuroprotection by drug treatment. Representative images of p-syn staining in SN sections in Control, SynFib, SynFib+Drug and Drug treatment groups at 16W. Scale bar = 500µm. **H.** Quantitative analysis of relative p-syn percentage signal intensity in the SN depicts significant reduction in p-syn following drug treatment. **I.** Representative SN sections co-stained for IBA1, p-syn and TH in Control, SynFib, SynFib+Drug and Drug treatment groups at 16W. Scale bar = 500µm. **J.** Quantitative analysis of percentage of area occupied by IBA1 depicts significant increase in injected side of SynFib group compared to Control. Further, a significant reduction in the percentage area of IBA1in the injected side of SynFib+Drug group compared to SynFib was observed. There was no significant increase in injected sides of SynFib+Drug compared to the control. **K.** Representative SN sections co-stained for GFAP, p-syn and TH in Control, SynFib, SynFib+Drug and Drug treatment groups at 16W. Scale bar = 500µm. **L.** Quantitative analysis of percentage area occupied by GFAP signal revealed a significant increase in the injected side of SynFib group relative to Control group which was significantly reduced in SynFib+Drug group depicting a reduction in the GFAP activation. Data are represented as mean ± SEM. For all the graphs D, F, H, J and L no of mice: Control (n=6), SynFib (n=7), SynFib+Drug (n=7), Drug (n=6). Ordinary one-way ANOVA, with Tukey’s multiple comparison test as the post hoc analysis for graphs D, F and H. (* P <0.05, ** P <0.01, *** P<0.001). Two-way ANOVA with Tukey’s multiple comparision test as post hoc for analyzing graphs J and L (* comparison between injected sides across groups; * P<0.05, ** P <0.01, *** P <0.001) (# comparison between uninjected and injected sides; ## P<0.01, ### P <0.001) Data points in red represent female mice; black represents male mice.

In comparison to the control, the SynFib groups showed a significant increase in the area occupied by activated IBA1 positive microglia (Fig 5I and J) that was significantly attenuated in the SynFib+ PD180970 group. Similarly, PD180970 treatment also caused a significant reduction in the area occupied by activated GFAP positive astrocytes in comparison to the SynFib group (Fig 5K and L).

## Discussion

The SynFib mouse model effectively reproduces key features of PD including progressive neurodegeneration, α-synuclein aggregation, and motor deficits, making it a highly valuable tool for studying disease mechanisms and therapeutic interventions.

Individual mice in SynFib model exhibited variability in cell loss reflecting the heterogeneous progression of symptom. This variability aligns with clinical findings, where the pace of neurodegeneration and symptom onset varies significantly among individuals. The motor deficits in SynFib model did not manifest with lesser cell loss at earlier timepoints, mirroring human PD, where motor defects become evident only after severe loss of TH innervations in the striatum (42, 43). During PD, as physiological stressors accumulate over prolonged periods, surviving neurons likely compensate for their function (23) by increasing their firing activity (44, 45) and enhancing dopamine release. These compensatory responses may explain the delayed onset of motor deficits in both PD patients and mouse models.

One of the key strengths of the SynFib model is its ability to reproduce α-synuclein pathology, including the formation of ThS-positive aggregates and the spread of p-syn to synaptically connected brain regions in striatum and cortex. This aligns with the Braak’s hypothesis which posits that aggregates propagate to more brain regions as the disease progresses. Lewy bodies, a hallmark of PD, are proteinase-K-resistant aggregates that stain with Thioflavin-S (ThS) in post-mortem human tissue (46). In the SynFib model, we observed similar ThS-positive aggregates, confirming amyloid pathology. Notably, some ThS-positive aggregates were negative for p-syn, suggesting that phosphorylation does not capture all aggregation states of α-synuclein in PD. We observed p-syn aggregates in MSNs of the striatum without significant cell loss, highlighting other neurons might possess mechanisms to clear the aggregates that DA neurons lack. These findings demonstrate that the model captures the higher vulnerability of DA neurons to degeneration, closely mimicking the human condition. Neurons attempted to clear aggregates via UPS, as shown by ubiquitin puncta colocalizing with p-syn. However, this proved insufficient to halt neurodegeneration, mirroring the downregulation of UPS in human PD(47). These findings highlight the SynFib model’s utility in studying PD-associated cellular dysfunctions observed in human brains.

We observed p-syn-positive structures in microglia on both the contralateral and injected sides, supporting their active involvement in α-synuclein propagation. The SynFib model showed increased activation of microglia and astrocytes, with peak IBA1 and GFAP staining at early timepoints. While surgery-induced inflammation could contribute, control groups undergoing surgery did not display comparable neuroinflammation, indicating neurodegeneration and α-synuclein aggregation as primary drivers of glial cell activation. Such observations have been made in the rat SynFib model as well where involvement of microglia in the transfer of aggregates was demonstrated (48). Microglia in the striatum mediated the transfer of aggregates from host DA neurons to grafted human embryonic stem cell region (48). In turn, these chronically activated microglial cells phagocytose the remaining neurons, thereby propagating neurodegeneration. While we did not observe many p-syn aggregates in astrocytes, their activation was evident in the SynFib model. Astrocytes are known to rapidly scavenge and degrade α-synuclein aggregates, making p-syn detection inside them challenging (49). However, activated astrocytes cannot release the neurotrophic factors required for neuronal maintenance, further contributing to neuronal death.

Our evaluation of PD180970, a c-Abl inhibitor, demonstrates the utility of the SynFib model for testing early-stage interventions. Chronic administration of PD180970 significantly reduced DA neuron loss and p-syn aggregation. Further, the drug also decreased neuroinflammation observed in SynFib model. These findings suggest that inhibition of α-synuclein aggregation and reduction of neuroinflammation serve as key mechanism for neuroprotection. Future studies should explore the underlying causes of sex differences in PD development and their implications for drug development. This study establishes proof of principle for the direct utility of the mouse SynFib model in testing early-stage interventions for PD, emphasizing the need for early disease detection.

The progressive nature of the SynFib model enables the investigation of all stages of PD, from early cellular dysfunction to late-stage pathology, highlights its potential for identifying biomarkers of disease onset. The SynFib mouse model holds promise for advancing our understanding of PD and accelerating the development of effective therapies.

## Materials and Methods

### Animals

Eight- to twelve-week-old C57BL/6J male and female mice were housed in the animal facility of the Indian Institute of Science Education and Research Thiruvananthapuram (IISER-TVM). The mice were maintained in individually ventilated cages with ad libitum access to food and water under a 12-hour light-dark cycle. All experimental procedures were approved by the Institutional Animal Ethics Committee (IAEC) and conducted in accordance with the guidelines established by the Committee for the Purpose of Control and Supervision of Experiments on Animals (CPCSEA), Government of India.

In the first part of the study, mice were unilaterally injected with AAV6-GFP (GFP group) overexpressing e-GFP (6 × 10¹⁰ gc/site in medial and lateral SN; n = 5), AAV6-α-synuclein (Syn group) overexpressing human α-synuclein (6 × 10¹⁰ gc/site in medial and lateral SN; n = 5), α-synuclein PFFs (Fib group) (2.5 µg/site in medial and lateral SN; n = 4), or a combination of AAV6-α-synuclein and α-synuclein pre-formed fibrils (SynFib group) (2.5 µg/site of PFFs and 6 × 10¹⁰ gc/site of AAV6-α-synuclein in medial and lateral SN; n = 10). A 1 µL total volume of each material was injected at the following coordinates: medial SN (AP = −3.2; DV = −4.6; ML = −1.1) and lateral SN (AP = −3.2; DV = −4.15; ML = −1.7). Behavioral and IHC analysis was done after 4W (Fig 1A).

In the second part of the study, the SynFib group was evaluated at two additional timepoints: 12W (n = 11) and 24W (n = 8). For comparison across timepoints, we included the mice from the 4W SynFib group (n = 10) from the first part of the study. Behavioral and IHC analyses were conducted at 12W and 24W (Fig 1F).

In the third part of the study, we established another cohort of mice for 16W, divided into four groups: the control group (unilaterally injected with DPBS in medial and lateral SN), the SynFib group, SynFib + PD180970 (SynFib + Drug), and PD180970 alone (Drug). Drug administration (5 mg/kg, i.p.) began one-week post-surgery and continued daily until 16W. All other groups received vehicle injections (1.875% DMSO/DPBS) to control for potential effects of injections or vehicle toxicity (Fig 5A).

### Statistical analysis

All the statistical analysis was performed using the GraphPad Prism software (9.3.1) Detailed materials and methods are provided in the supplementary information file S1.

## Acknowledgements

This work was supported by the DBT/Wellcome Trust India Alliance Fellowship (IA/E/17/1/503664) awarded to Poonam Thakur. Funding support from SERB (SRG/2021/000981) to Poonam Thakur is also acknowledged. Santhosh Kumar Subramanya is supported by a fellowship from Department of Biotechnology, Government of India. We are grateful to Malin Parmer for generous gift of the AAV-α-synuclein. We thank Akshaya Rajan, Unnati Agrawal and Jyotirmay Srivastava for their support with certain mice experiments and imaging. We acknowledge the assistance from Gargi Dutta, Sharadha Sheshadri and Naveen Balachandran for behavioural analysis. We also thank Dr. Abhilasha Joshi for suggestions regarding gait analysis.

## References

1. P. P. Michel, E. C. Hirsch, S. Hunot, Understanding Dopaminergic Cell Death Pathways in Parkinson Disease. Neuron 90, 675–691 (2016).

2. N. Panicker, P. Ge, V. L. Dawson, T. M. Dawson, The cell biology of Parkinson’s disease. J Cell Biol 220, e202012095 (2021).

3. J. Nam, C. T. Richie, B. K. Harvey, M. H. Voutilainen, Delivery of CDNF by AAV-mediated gene transfer protects dopamine neurons and regulates ER stress and inflammation in an acute MPTP mouse model of Parkinson’s disease. Sci Rep 14, 16487 (2024).

4. Q. Li, et al., Partial depletion and repopulation of microglia have different effects in the acute MPTP mouse model of Parkinson’s disease. Cell Proliferation 54, e13094 (2021).

5. M. Santoro, et al., Neurochemical, histological, and behavioral profiling of the acute, sub-acute, and chronic MPTP mouse model of Parkinson’s disease. Journal of Neurochemistry 164, 121–142 (2023).

6. M. Santoro, et al., Mapping of catecholaminergic denervation, neurodegeneration, and inflammation in 6-OHDA-treated Parkinson’s disease mice. Res Sq rs.3.rs-5206046 (2024). 10.21203/rs.3.rs-5206046/v1.

7. S. H. Moon, Y. Kwon, Y. E. Huh, H. J. Choi, Trehalose ameliorates prodromal non-motor deficits and aberrant protein accumulation in a rotenone-induced mouse model of Parkinson’s disease. Arch. Pharm. Res. 45, 417–432 (2022).

8. L. Wang, et al., Astaxanthin ameliorates dopaminergic neuron damage in paraquat-induced SH-SY5Y cells and mouse models of Parkinson’s disease. Brain Research Bulletin 202, 110762 (2023).

9. A. Slézia, et al., Behavioral, neural and ultrastructural alterations in a graded-dose 6-OHDA mouse model of early-stage Parkinson’s disease. Sci Rep 13, 19478 (2023).

10. R. Lal, A. singh, S. watts, K. Chopra, Experimental models of Parkinson’s disease: Challenges and Opportunities. European Journal of Pharmacology 980, 176819 (2024).

11. M. El-Gamal, et al., Neurotoxin-Induced Rodent Models of Parkinson’s Disease: Benefits and Drawbacks. Neurotox Res 39, 897–923 (2021).

12. L. Glasl, et al., Pink1-deficiency in mice impairs gait, olfaction and serotonergic innervation of the olfactory bulb. Experimental Neurology 235, 214–227 (2012).

13. P. W.-L. Ho, et al., Age-dependent accumulation of oligomeric SNCA/α-synuclein from impaired degradation in mutant LRRK2 knockin mouse model of Parkinson disease: role for therapeutic activation of chaperone-mediated autophagy (CMA). Autophagy 16, 347–370 (2020).

14. H. Liu, et al., Aberrant mitochondrial morphology and function associated with impaired mitophagy and DNM1L-MAPK/ERK signaling are found in aged mutant Parkinsonian LRRK2R1441G mice. Autophagy 17, 3196–3220 (2021).

15. Y. Xiong, et al., Robust kinase- and age-dependent dopaminergic and norepinephrine neurodegeneration in LRRK2 G2019S transgenic mice. Proceedings of the National Academy of Sciences 115, 1635–1640 (2018).

16. M. Wegrzynowicz, et al., Depopulation of dense α-synuclein aggregates is associated with rescue of dopamine neuron dysfunction and death in a new Parkinson’s disease model. Acta Neuropathol 138, 575–595 (2019).

17. H.-W. Kim, et al., Genetic reduction of mitochondrial complex I function does not lead to loss of dopamine neurons in vivo. Neurobiology of Aging 36, 2617–2627 (2015).

18. K. Mao, et al., Poly (ADP-ribose) polymerase 1 inhibition prevents neurodegeneration and promotes α-synuclein degradation via transcription factor EB-dependent autophagy in mutant α-synucleinA53T model of Parkinson’s disease. Aging Cell 19, e13163 (2020).

19. M. Decressac, B. Mattsson, M. Lundblad, P. Weikop, A. Björklund, Progressive neurodegenerative and behavioural changes induced by AAV-mediated overexpression of α-synuclein in midbrain dopamine neurons. Neurobiology of Disease 45, 939–953 (2012).

20. A. Ulusoy, M. Decressac, D. Kirik, A. Björklund, Viral vector-mediated overexpression of α-synuclein as a progressive model of Parkinson’s disease. Prog Brain Res 184, 89–111 (2010).

21. C. W. Ip, et al., AAV1/2-induced overexpression of A53T-α-synuclein in the substantia nigra results in degeneration of the nigrostriatal system with Lewy-like pathology and motor impairment: a new mouse model for Parkinson’s disease. Acta Neuropathologica Communications 5, 11 (2017).

22. M. Negrini, et al., Sequential or Simultaneous Injection of Preformed Fibrils and AAV Overexpression of Alpha-Synuclein Are Equipotent in Producing Relevant Pathology and Behavioral Deficits. Journal of Parkinson’s Disease 12, 1133–1153 (2022).

23. K. Albert, et al., Downregulation of tyrosine hydroxylase phenotype after AAV injection above substantia nigra: Caution in experimental models of Parkinson’s disease. Journal of Neuroscience Research 97, 346–361 (2019).

24. K. C. Luk, et al., Pathological α-Synuclein Transmission Initiates Parkinson-like Neurodegeneration in Nontransgenic Mice. Science 338, 949–953 (2012).

25. M. Masuda-Suzukake, et al., Prion-like spreading of pathological α-synuclein in brain. Brain 136, 1128–1138 (2013).

26. L. A. Volpicelli-Daley, K. C. Luk, V. M.-Y. Lee, Addition of exogenous α-synuclein preformed fibrils to primary neuronal cultures to seed recruitment of endogenous α-synuclein to Lewy body and Lewy neurite–like aggregates. Nat Protoc 9, 2135–2146 (2014).

27. N. K. Polinski, A Summary of Phenotypes Observed in the In Vivo Rodent Alpha-Synuclein Preformed Fibril Model. Journal of Parkinson’s Disease 11, 1555–1567 (2021).

28. P. Thakur, et al., Modeling Parkinson’s disease pathology by combination of fibril seeds and α-synuclein overexpression in the rat brain. Proceedings of the National Academy of Sciences 114, E8284–E8293 (2017).

29. A. Björklund, F. Nilsson, B. Mattsson, D. B. Hoban, M. Parmar, A Combined α-Synuclein/Fibril (SynFib) Model of Parkinson-Like Synucleinopathy Targeting the Nigrostriatal Dopamine System. Journal of Parkinson’s Disease 12, 2307–2320 (2022).

30. F. Fredlund, et al., Ciita Regulates Local and Systemic Immune Responses in a Combined rAAV-α-synuclein and Preformed Fibril-Induced Rat Model for Parkinson’s Disease. J Parkinsons Dis 14, 693–711 (2024).

31. C. J. Barnum, et al., Peripheral administration of the selective inhibitor of soluble tumor necrosis factor (TNF) XPro®1595 attenuates nigral cell loss and glial activation in 6-OHDA hemiparkinsonian rats. J Parkinsons Dis 4, 349–360 (2014).

32. J. F. Dorsey, R. Jove, A. J. Kraker, J. Wu, The pyrido[2,3-d]pyrimidine derivative PD180970 inhibits p210Bcr-Abl tyrosine kinase and induces apoptosis of K562 leukemic cells. Cancer Res 60, 3127–3131 (2000).

33. A.-L. Mahul-Mellier, et al., c-Abl phosphorylates α-synuclein and regulates its degradation: implication for α-synuclein clearance and contribution to the pathogenesis of Parkinson’s disease. Human Molecular Genetics 23, 2858–2879 (2014).

34. Z.-H. Zhou, Y.-F. Wu, X. Wang, Y.-Z. Han, The c-Abl inhibitor in Parkinson disease. Neurol Sci 38, 547–552 (2017).

35. H. S. Ko, et al., Phosphorylation by the c-Abl protein tyrosine kinase inhibits parkin’s ubiquitination and protective function. Proc Natl Acad Sci U S A 107, 16691–16696 (2010).

36. R. Wu, et al., c-Abl–p38α signaling plays an important role in MPTP-induced neuronal death. Cell Death Differ 23, 542–552 (2016).

37. S. Brahmachari, et al., Activation of tyrosine kinase c-Abl contributes to **α**-synuclein–induced neurodegeneration. J Clin Invest 126, 2970–2988 (2016).

38. S. Sn, et al., Small molecule modulator of aggrephagy regulates neuroinflammation to curb pathogenesis of neurodegeneration. EBioMedicine 50, 260–273 (2019).

39. J.-Y. Shin, et al., Dual inhibition of aminoacyl-tRNA synthetase interacting multifunctional protein-2 and α-synuclein by steroid derivative is neuroprotective in Parkinson’s model. iScience 27, 111165 (2024).

40. S. S. Karuppagounder, et al., The c-Abl inhibitor IkT-148009 suppresses neurodegeneration in mouse models of heritable and sporadic Parkinson’s disease. Science Translational Medicine 15, eabp9352 (2023).

41. M. Zhong, Y. Wang, G. Lin, F.-F. Liao, F.-M. Zhou, Dopamine-independent development and maintenance of mouse striatal medium spiny neuron dendritic spines. Neurobiology of Disease 181, 106096 (2023).

42. G. M. Halliday, H. McCann, The progression of pathology in Parkinson’s disease. Annals of the New York Academy of Sciences 1184, 188–195 (2010).

43. R. Yilmaz, F. Hopfner, T. van Eimeren, D. Berg, Biomarkers of Parkinson’s disease: 20 years later. J Neural Transm 126, 803–813 (2019).

44. A. Tozzi, et al., Dopamine-dependent early synaptic and motor dysfunctions induced by α-synuclein in the nigrostriatal circuit. Brain 144, 3477–3491 (2021).

45. A. Dagra, et al., α-Synuclein-induced dysregulation of neuronal activity contributes to murine dopamine neuron vulnerability. npj Parkinsons Dis. 7, 1–22 (2021).

46. K. Tanji, et al., Proteinase K-resistant α-synuclein is deposited in presynapses in human Lewy body disease and A53T α-synuclein transgenic mice. Acta Neuropathol 120, 145–154 (2010).

47. B. Dehay, et al., Pathogenic Lysosomal Depletion in Parkinson’s Disease. J Neurosci 30, 12535–12544 (2010).

48. D. B. Hoban, et al., Impact of α-synuclein pathology on transplanted hESC-derived dopaminergic neurons in a humanized α-synuclein rat model of PD. Proc Natl Acad Sci U S A 117, 15209–15220 (2020).

49. Y. Yang, et al., Therapeutic functions of astrocytes to treat α-synuclein pathology in Parkinson’s disease. Proceedings of the National Academy of Sciences 119, e2110746119 (2022).

