## Supplementary material for "A Novel Mouse Model of Parkinson’s Disease for Investigating Progressive Pathology and Neuroprotection"

**This PDF file includes:**

Supplementary text
Figures S1 to S7
SI References

### Supplementary Data

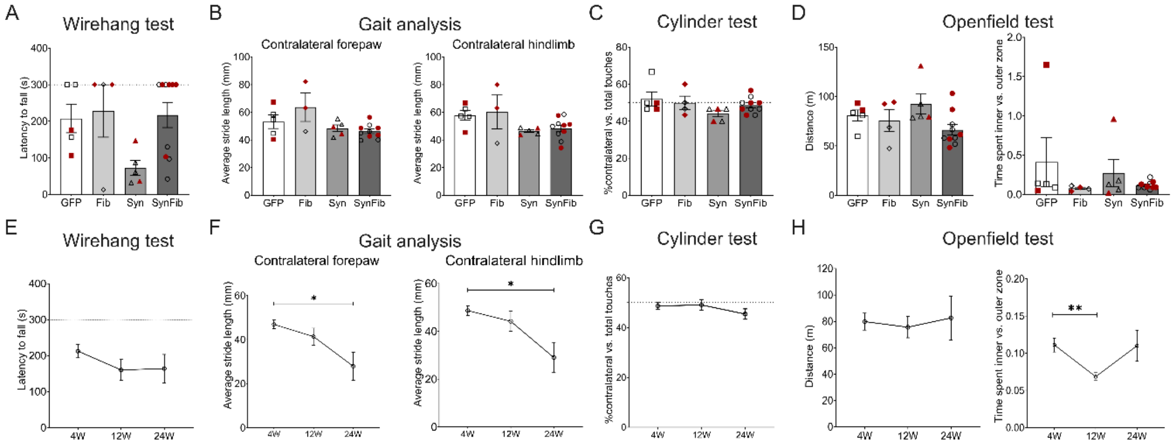


***Fig S1: Behavioral Assessment of motor function in mice***

***A.*** *Wire hang test for neuromuscular strength showed no significant differences among groups. No of mice: GFP (n = 5), Fib (n = 4), Syn (n = 5), SynFib (n = 10)* ***B.*** *Gait analysis revealed no significant differences in average stride length among groups. No of mice: GFP (n = 5), Fib (n = 3), Syn (n = 5), SynFib (n = 10).* ***C.*** *Cylinder test showing a similar percentage of contralateral paw usage relative to the total number of paw touches across groups at 4W. No of mice: GFP (n = 5), Fib (n = 4), Syn (n = 5), SynFib (n = 10).* ***D.*** *Open field test at 4W in GFP, Fib, Syn, and SynFib groups, indicating no significant changes in locomotor activity. No of mice:-: GFP (n = 5), Fib (n = 4), Syn (n = 5), SynFib group (n = 10).* ***E*** *Wire hang test for neuromuscular strength displayed a non-significant decrease in latency to fall over time in the SynFib group.* ***F.*** *Gait analysis showed a significant reduction in average stride length at 24W compared to 4W. No of mice for E and F: 4W (n = 29), 12W (n = 19), 24W (n = 8).* ***G.*** *Cylinder test revealing a non-significant decrease in contralateral paw usage over time in the SynFib group. No of mice: 4W (n = 29), 12W (n = 19), 24W (n = 8).* ***H.*** *Open field test revealed no change in the total distance travelled by SynFib mice over time. A significant reduction in time spent in the inner vs outer zone was observed at 12W w.r.t. 4W. No of mice: 4W (n = 29), 12W (n = 19), 24W (n = 8). All data are represented as mean ± SEM. Statistical analysis was performed using one-way ANOVA followed by Kruskal-Wallis multiple comparison post hoc test (* P<0.05, **P < 0.01). Data points in red represent female mice; black represents male mice.*

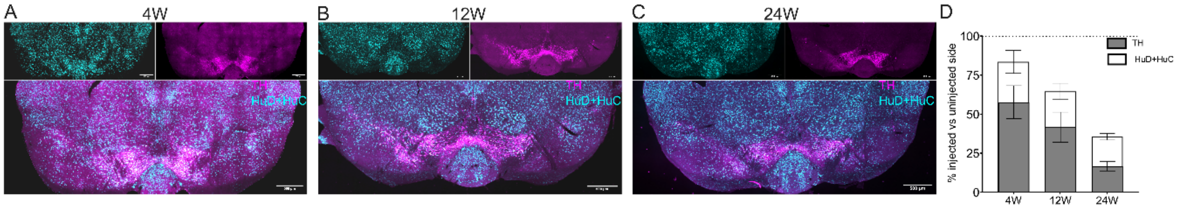


***Fig S2. Progressive loss of neurons in the SN.***
***A-C****. Representative images of SN sections stained for HuD+HuC and TH at 4, 12 and 24W respectively. (Scale bar = 500µm)* ***D.*** *Quantification of TH-positive neurons relative to HuD+HuC on the injected side revealed a time-dependent reduction in both TH-positive and HuD+HuC-stained cells. The greater extent of loss in TH-positive neurons compared to the loss of HuD+HuC-stained cells suggests TH downregulation rather than a corresponding loss of general neurons in*

the SN. No of mice: 4W (n=8), 12W (n=11), 24W (n=8). All data are represented as mean ± SEM. Statistical analysis was performed using one-way ANOVA followed by Kruskal-Wallis multiple comparison post hoc test applied separately for TH and HuD+HuC cell counts.

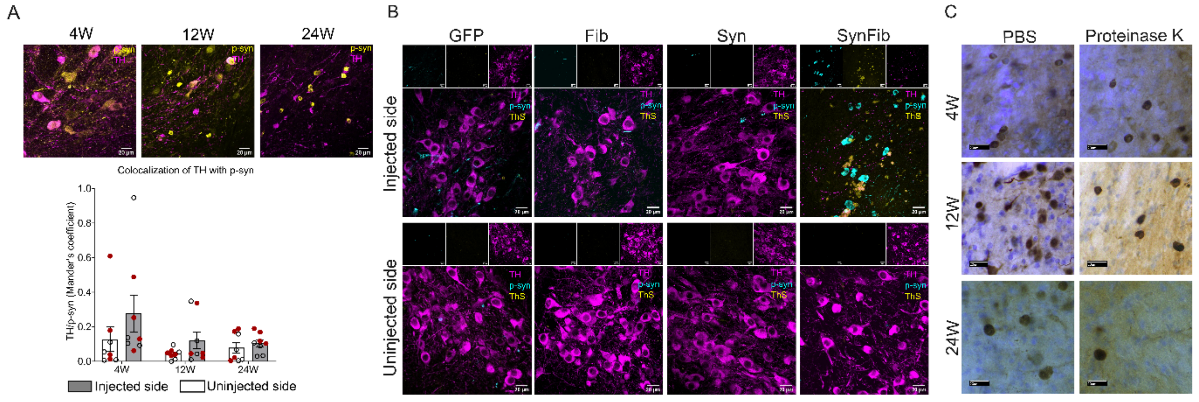


***Fig S3: Lewy-like characteristics of aggregates in the SynFib group***
***A.*** *High magnification images of sections co-stained with TH and p-syn at 4, 12 and 24W show higher colocalization of TH with p-syn in the injected side compared to the uninjected side that reduced by 24W due to progressive loss of TH-positive cells. No of mice 4W (n=8), 12W(n=8), 24W (n=8). Scale bars = 20µm. All data are represented as mean ± SEM. Statistical analysis was performed using two-way ANOVA with Tukey's multiple comparision test as post hoc. Data points in red represent female mice; black represents male mice.* ***B.*** *Representative SN sections co-stained for p-syn, ThS and TH in GFP, Fib, Syn, and SynFib groups at 4W. Only SynFib groups showed ThS positive aggregates. Scale bars = 20µm.* ***C.*** *Representative SN sections from SynFib mice at 4, 12 and 24W stained for p-syn, showing resistance to proteinase-K digestion. Scale bar = 20µm.*

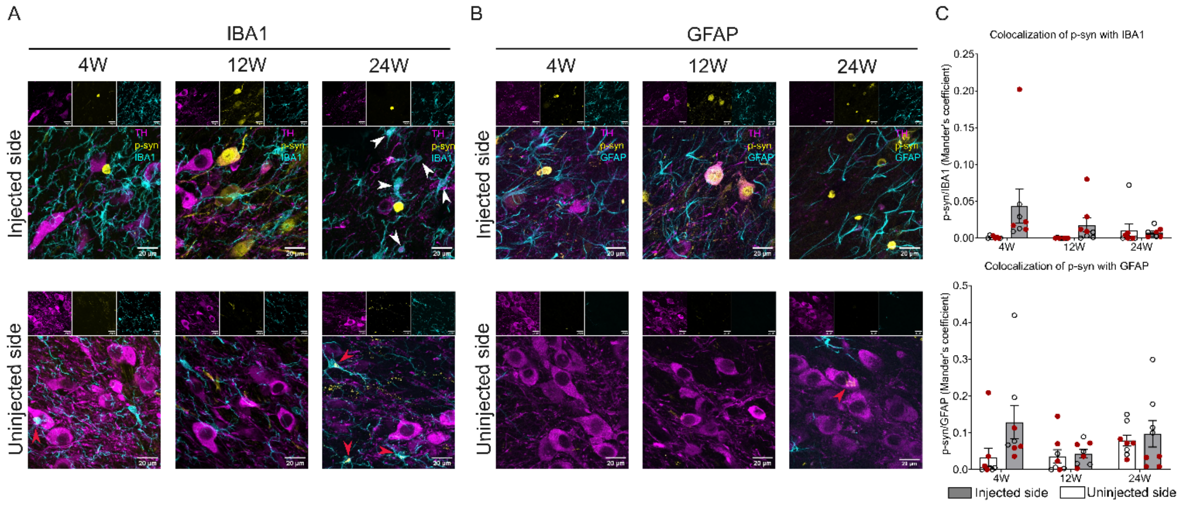


***Figure S4: p-syn colocalizes with both microglia and astrocytes and reduces over time***
***A.*** *Representative high-magnification SN sections co-stained for IBA1, p-syn and TH in the SynFib group at 4W, 12W, and 24W. Microglia engulfing the TH-positive debris were observed indicating their role in mediating neurodegeneration (white arrowheads). Contralateral spread of p-syn in the microglia can be visualized (red arrowheads). Scale bar = 20µm.* ***B.*** *Representative high-magnification SN sections co-stained for GFAP, p-syn and TH in the SynFib group at 4W, 12W, and 24W. Scale bar = 20µm. Contralateral spread of p-syn can be visualized (red arrowheads).* ***C.*** *Colocalization analysis revealed modest increase in colocalization of p-syn with both microglia*

*and astrocytes in the injected side compared to the uninjected side at early timepoint which reduced over time. No of mice for IBA1: 4W (n=8), 12W(n=8), 24W (n=8); No of mice for GFAP: 4W (n=8), 12W(n=8), 24W (n=8). All data are represented as mean ± SEM. Statistical analysis was performed using two-way ANOVA was used for the analysis with Tukey's multiple comparison as post-hoc. Data points in red represent female mice; black represents male mice*

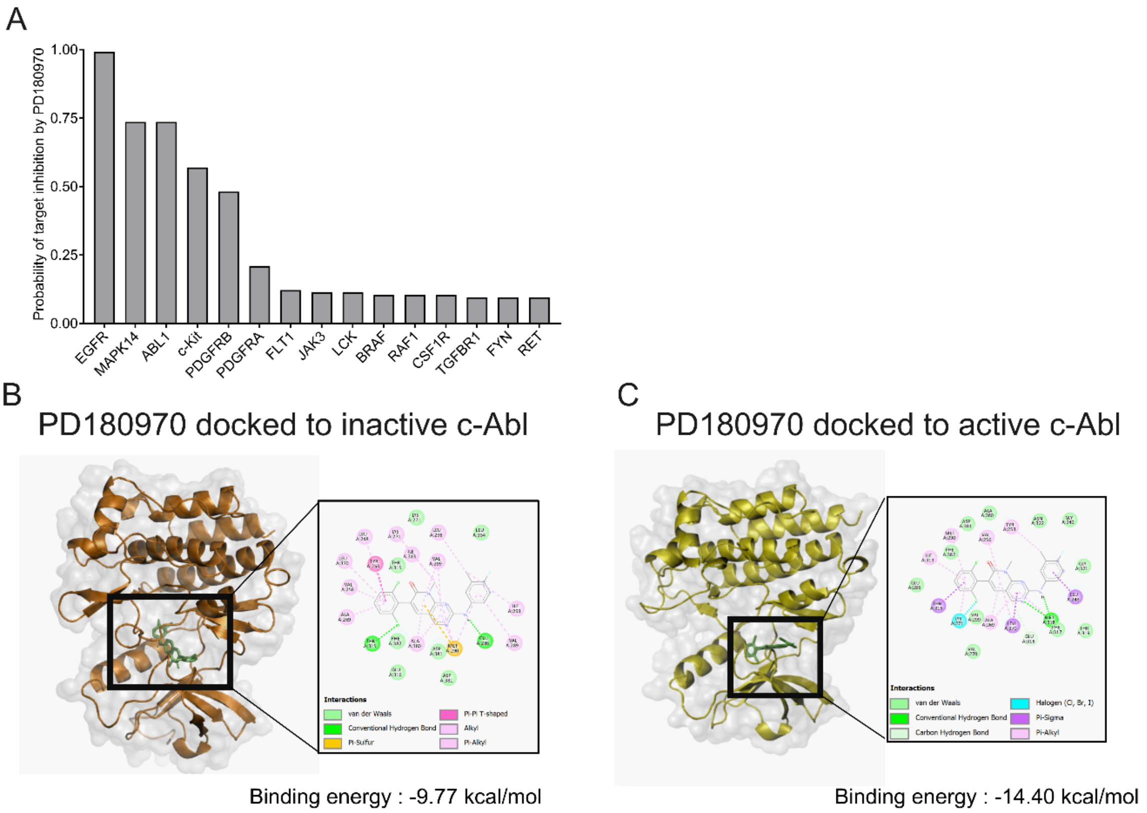


***Fig S5: Molecular docking of PD180970 with c-Abl and target prediction***
***A.*** *Potential target interactions of PD180970, as predicted by Swiss Target Prediction, identified mouse c-Abl among the top three predicted targets.* ***B and C.*** *3-dimensional and 2-dimensional representation of molecular interactions of PD180970 with the inactive c-Abl conformation (PDB ID: 1IEP) and the active c-Abl conformation (PDB ID: 1M52), respectively, suggesting that PD180970 can inhibit both conformations.*

#### Materials and methods

##### Stereotaxic injection material

PFFs were generated in the lab of Kelvin Luk using full-length human-α-synuclein using the methodology previously described (1). Characterization of PFFs was carried out at IISER-TVM. Briefly, recombinant α-synuclein was expressed in the BL21 cells under an inducible promoter. IPTG was used to induce the expression, and the cells were harvested, lysed, and α-synuclein was purified using gel filtration and ion-exchange chromatographic methods. The aggregates were generated *in-vitro* using monomeric α-synuclein incubated at 37°C with constant agitation of 1000 rpm in tris-buffered saline.

For the Thioflavin-T fluorescence assay, a final concentration of 3.5µM of α-synuclein was mixed with 10µM of Thioflavin T. Fluorescence measurements were carried out in a 96-well dark plate in triplicates using TECAN Infinite M200 Pro at an excitation of 440 nm and emission of 480 nm at 37°C. PFF exhibited higher fluorescence intensity in the presence of Thioflavin T (ThT) dye compared to monomeric α-synuclein (Fig S6A).

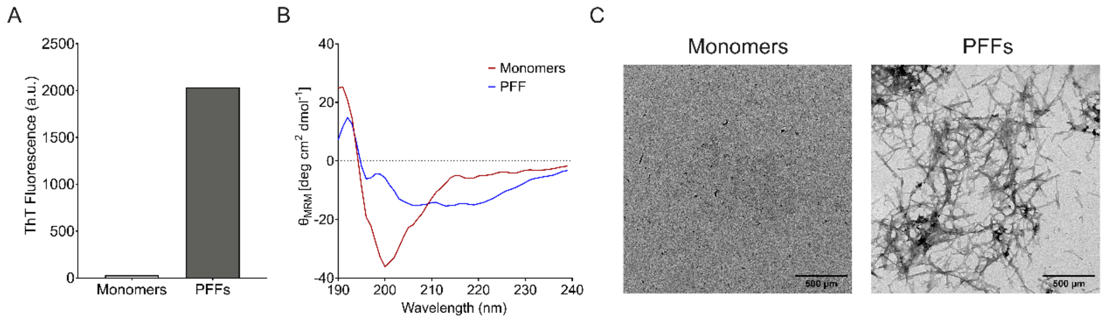


***Fig S6: Characterization of α-synuclein pre-formed fibrils reveals fibrillar structure***

***A.*** *ThT fluorescence readings of α-synuclein monomers and PFFs indicating the presence of amyloid aggregates in PFFs.* ***B.*** *CD spectroscopy of α-synuclein monomers and PFFs demonstrating the characteristic random coil structure in monomers and β-sheet structure of PFFs.* ***C.*** *Transmission electron micrographs of monomers and PFFs confirming the fibrillar morphology of PFFs. Scale bar = 500 µm .*

To obtain the CD spectra, α-synuclein was mixed with PBS to a final concentration of 20mM. Far-UV CD spectra was recorded in BioLogic MOS-500 CD Spectrometer using 0.1 cm cuvette at 25ºC in the 190-240 nm range with a 1 nm step reading at 1 nm/sec. Aggregation was confirmed using circular dichroism (CD) spectroscopy, with PFF displaying a characteristic dip at 220 nm, indicative of a β-sheet structure (Fig S6B). In contrast, monomeric α-synuclein showed a dip at 190 nm, consistent with a random coil conformation.

Transmission electron microscopy (TEM) was performed by blotting PFFs and monomers on the carbon grids (Type-B, Ted Pella, Inc. 10814-F), negatively stained with uranyl acetate and visualized using FEI-TECNAI-G2 Spirit BioTwin Transmission Electron Microscope. TEM imaging revealed multiple filamentous structures in PFF, while no such structures were observed in the monomeric form (Fig S6C).

AAV2/6 vectors used to overexpress human-α-synuclein or GFP under the synapsin-1 promoter were generated at Lund University Viral Vector Core. The expression of α-synuclein in the Syn and SynFib groups was validated through IHC for human α-synuclein using the syn211 antibody (Fig S7). Similarly, GFP expression was validated by immunostaining in SN and STR sections. Expression of both α-synuclein and GFP was mainly restricted to the injected side of the brain.

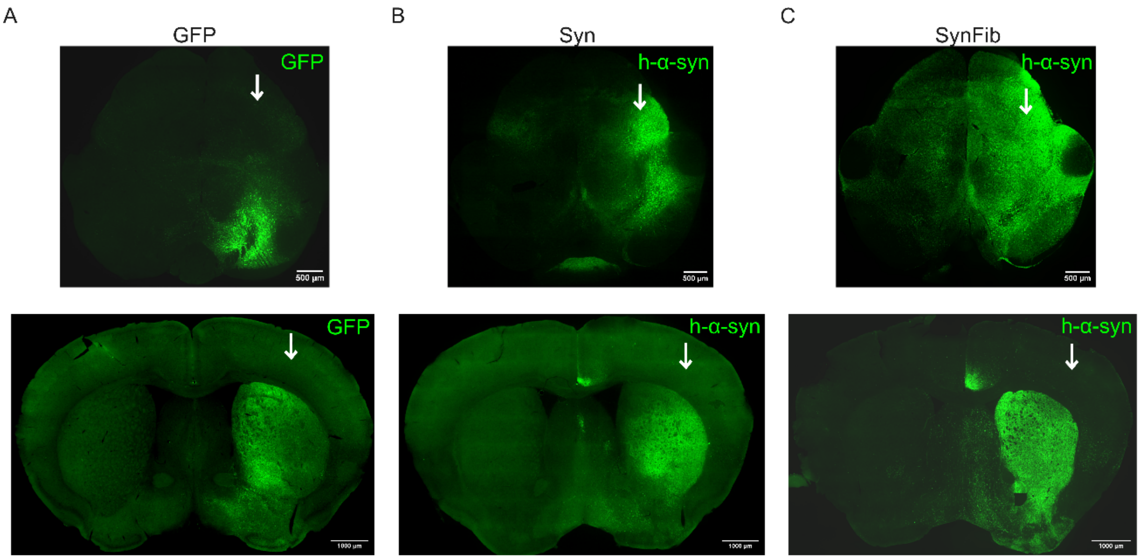


***Fig S7: Unilateral AAV injection in the SN leads to overexpression in the nigro-striatal pathway.***
*Representative images of SN and STR showing unilateral overexpression of* ***A.*** *GFP in the GFP group* ***B.*** *human α-synuclein in the Syn group* ***C.*** *human α-synuclein in the SynFib group. Scale bar, SN = 500µm and STR = 1000µm.*

#### Stereotaxic surgeries

Surgeries were performed after mice were anaesthetised using isoflurane. Prior to injection, the fibrils were sonicated for 8 minutes using a Power Sonic 410 bath sonicator, with a 30-second on and 30-second off pulse cycle. Hamilton syringe (CAL #87930 26G/51mm/3) was used for infusion at the rate of 250nL/min using a Kd Scientific 100 syringe pump. After the injection, the syringe was left in place for an additional 5 minutes, and then slowly retracted. Afterwards, the skin was sutured, and antibiotic powder was applied at the site of the incision. The animals were kept on an electrically heated pad until their full movement recovered. Post-surgery, mice were closely monitored for a week with regular weight measurements.

#### Administration of PD180970

For administration into mouse, the drug was dissolved in 100% DMSO at a concentration of 40mg/ml. This stock solution was further diluted in DPBS to make a concentration of 0.75mg/ml corresponding to a concentration of 5mg/kg body weight of injections taking the reference of a 200 µL injection volume for a 30 gm mouse.

#### Behavioural tests

Behavioral assessments were conducted every 4 weeks until the 24W timepoint (except for the drug cohort, which was assessed until 16W). All animals were acclimated to the experimental room for one hour prior to behavioral testing. Motor functions were evaluated using the open field test, gait analysis, cylinder test and wire hang test.

*Open field test*
Mice were allowed to explore a 50cm X 50cm open arena for 30 minutes (5 minutes acclimatization time + 25 minutes evaluation time) while being video recorded from the top. The recording was fed into AnyMaze software for continuous tracking of mouse movements in the field. Total distance travelled and time spent in the inner versus outer zone were analyzed.

*Gait analysis*
Mice were made to walk across a transparent perspex corridor with videos recorded from the bottom. Videos were preprocessed, and Visual Gait Lab (2) was used for extracting and labelling the frames and subsequently computing and analyzing the gait parameters. The average stride length of each paw was plotted and represented.

*Cylinder test*

Mice were placed inside a glass beaker (cylinder) and video recorded. The videos were played and the number of left paw touches (contralateral) versus total touches on the walls of the cylinder were manually scored in the recordings to get an estimate for forelimb asymmetry

*Wire hang test*

Mice were made to grip on a wire mesh and were gently shaken to induce grip. The mesh was inverted and placed on a stand and latency to fall in seconds was recorded using a handheld timer.

**Histology**

Mice were perfused transcardially with ice-cold PBS followed by 4% paraformaldehyde (PFA). The brains were removed and post-fixed in 4% PFA for 24 hours, followed by dehydration and storage in 20% sucrose at 4°C. The brains were sectioned using a vibratome (Leica VT1000 S) at a thickness of 30μm in series of 5 and preserved in PBS + 0.02% sodium azide until staining.

The slices were rinsed with PBS, followed by permeabilization with PBS+ 0.1% Triton X 100, antigen retrieval by boiling with citrate buffer at 80°C, blocking with 2% BSA, followed by primary antibody incubation for 12 hrs at 4°C and secondary antibody treatment for 6-8 hrs at 4°C. Some sections were co-stained with DAPI by incubation for 10 minutes at room temperature. For the TH and p-syn stained sections, the chromogenic development method based on DAB was preferred. Briefly, after secondary antibody (biotinylated) incubation, the sections were washed thrice in PBST. The development solution was prepared as per the manufacturer's recommendations (Vectastain ABC Kit, peroxidase PK-4000) and slices were incubated in ABC reagent for 30 min at room temperature. Following the incubation, the sections were washed twice and revealed with DAB (sigma DB001-1G) at 0.05%DAB in 0.01%H2O2 solution in PBS. Some sections were counterstained with 0.1% cresyl violet stain. The extent of DA neurodegeneration was evaluated using staining for TH in nigral and striatal slices. Cell count was performed in 6-8 slices throughout the rostro-caudal extent of the brain. The slices were mounted in rostral to caudal orientation and preserved under DPX mountant and imaged.

**Table 1.** List of primary antibodies.

| Target antigen | Brand | Catalog no. | Host | Dilution |
|---|---|---|---|---|
| **Tyrosine hydroxylase** | Abcam | ab112 | Rabbit | 1:700 |
| **Tyrosine hydroxylase** | Abcam | ab76442 | Chicken | 1:500 |
| **Human-α-synuclein (syn211)** | Abcam | ab80627 | Mouse | 1:1000 |
| **p-syn S129** | Abcam | ab184674 | Mouse | 1:8000 |
| **IBA1** | Abcam | ab178847 | Rabbit | 1:1000 |
| **GFAP** | Abcam | ab68428 | Rabbit | 1:1000 |
| **DARPP-32** | Abcam | ab40801 | Rabbit | 1:1000 |
| **Ubiquitin** | Abcam | ab7780 | Rabbit | 1:1000 |
| **HuD+HuC** | Abcam | ab184267 | Rabbit | 1:1000 |

**Table 2.** List of secondary antibodies

| Target species | Brand | Catalog no | Host | Dilution | Conjugate |
|---|---|---|---|---|---|
| **Rabbit** | Abcam | ab207995 | Goat | 1:500 | Biotin |
| **Mouse** | Abcam | ab6788 | Goat | 1:1000 | Biotin |
| **Rabbit** | Cell signaling technology | #4414 | Goat | 1:1000 | Alexa flour 647 |
| **Mouse** | Cell signaling technology | #4408 | Goat | 1:1000 | Alexa flour 488 |
| **Chicken** | Abcam | ab150176 | Goat | 1:1000 | Alexa flour 594 |

**Imaging and quantification**

DAB and Nissl-stained brain slices were imaged using a Radical (RXL-r4) brightfield microscope, with image acquisition performed using HS Analysis scan software. Low-magnification fluorescence images (20X) were acquired using an Olympus BX63 fluorescence microscope. High-magnification fluorescence images (60X and 100X) were captured using either an Olympus inverted confocal microscope (Fluoview FV3000) or a Leica upright confocal microscope (Stellaris 5). Quantification of parameters such as cell number and intensity in light microscope images was

performed using particle analyzer of ImageJ for cell counts and mean intensity for striatal TH density. For multichannel fluorescence images, individual channels were separated, and intensity measurements were taken in the marked nigral regions using ImageJ. Cell quantification in fluorescence images was conducted using QuPath software. For p-syn quantification in the PD180970 treated cohort, the IBA1 and GFAP images were split into channels and the mean fluorescence intensity of p-syn at SN was measured using ImageJ at both injected and uninjected sides. The average values were plotted as percantage injected side vs. uninjected side. The IBA1 and GFAP quantification was performed in ImageJ by calculating percentage area occupied as mentioned previously (3). Briefly, the multiple channels were split and SN region was marked in the uninjected side and copied and flipped on the injected side. Later, the 8-bit images were despeckled, thresholded, converted to binary and again despeckled and denoised removed noise. The percentage area occupied in the IBA1 and GFAP channels were measured and analyzed.
The colocalization analysis was performed using JaCoP plugin in the ImageJ as mentioned previously (4). The 20X images were split into individual channels and background was subtracted with rolling ball radius 0.50 using ImageJ. JaCoP analysis was performed and the Mander's coefficient in SN of both injected and un-injected side were used for colocalization analysis.

**Molecular docking and off-target prediction**

PD180970 was docked into the kinase domain of both active and in-active mouse c-Abl protein. The crystal structures were downloaded from RCSB-PDB as a pdb file (PDB ID 1IEP for inactive and 1M52 for active c-Abl). The proteins were prepared by removing the co-crystallized ligands and chain B using Autodock software (5). Both the proteins and ligand files were exported in pdbqt format. For docking, we used Autodock 4.2 software. We used prankweb (6) to determine the active site and the coordinates were used for the docking process. The Autogrid and Autodock programs were run. The docked confirmations were exported as ranks based on binding energies in pdb format. We used Discovery Studio software v24.1.0.23298 to visualize the ligand-protein interactions. The Swiss Target Prediction tool was used to predict the off-target binding proteins.

**Statistical analysis**

For the quantification of imaging data, mean intensity values, cell counts or percentage area occupied valued were quantified from ImageJ and statistical analysis was performed in GraphPad Prism software (9.3.1) that removed an outlier from the SynFib group in the p-syn quantification data. All the analyses in Fig 1, Fig S1, Fig 2B and Fig 3C were performed using non-parametric one-way ANOVA followed by Kruskal-Wallis multiple comparison post hoc test. Fig 2D, Fig 4C, 4D, 4G, 4H and 5B, 5J, 5L, S3A, and S4C were analyzed using two-way ANOVA using Tukey's multiple comparison as post hoc. Fig 5B, 5D, 5Fand 5H were analyzed using ordinary one-way ANOVA with Tukey's post hoc test.

**SI References:**


1. L. A. Volpicelli-Daley, *et al.*, Exogenous α-Synuclein Fibrils Induce Lewy Body Pathology Leading to Synaptic Dysfunction and Neuron Death. *Neuron* **72**, 57–71 (2011).

2. OSF | Visual Gait Lab: A user-friendly approach to gait analysis. Available at: https://osf.io/2ydzn/ [Accessed 6 January 2025].

3. Quantifying Microglia Morphology from Photomicrographs of Immunohistochemistry Prepared Tissue Using ImageJ. Available at: https://app.jove.comquantifying microglia morphology from photomicrographs of immunohistochemistry prepared tissue using imagej [Accessed 29 January 2025].

4. S. Bido, *et al.*, Microglia-specific overexpression of α-synuclein leads to severe dopaminergic neurodegeneration by phagocytic exhaustion and oxidative toxicity. *Nat Commun* **12**, 6237 (2021).

5. G. M. Morris, D. S. Goodsell, R. Huey, A. J. Olson, Distributed automated docking of flexible ligands to proteins: Parallel applications of AutoDock 2.4. *J Computer-Aided Mol Des* **10**, 293–304 (1996).

6. PrankWeb: a web server for ligand binding site prediction and visualization | Nucleic Acids Research | Oxford Academic. Available at: https://academic.oup.com/nar/article/47/W1/W345/5494740 [Accessed 7 January 2025].
